## Supplementary Movies for "Active state structures of a bistable visual opsin bound to G proteins"

### Slide 1
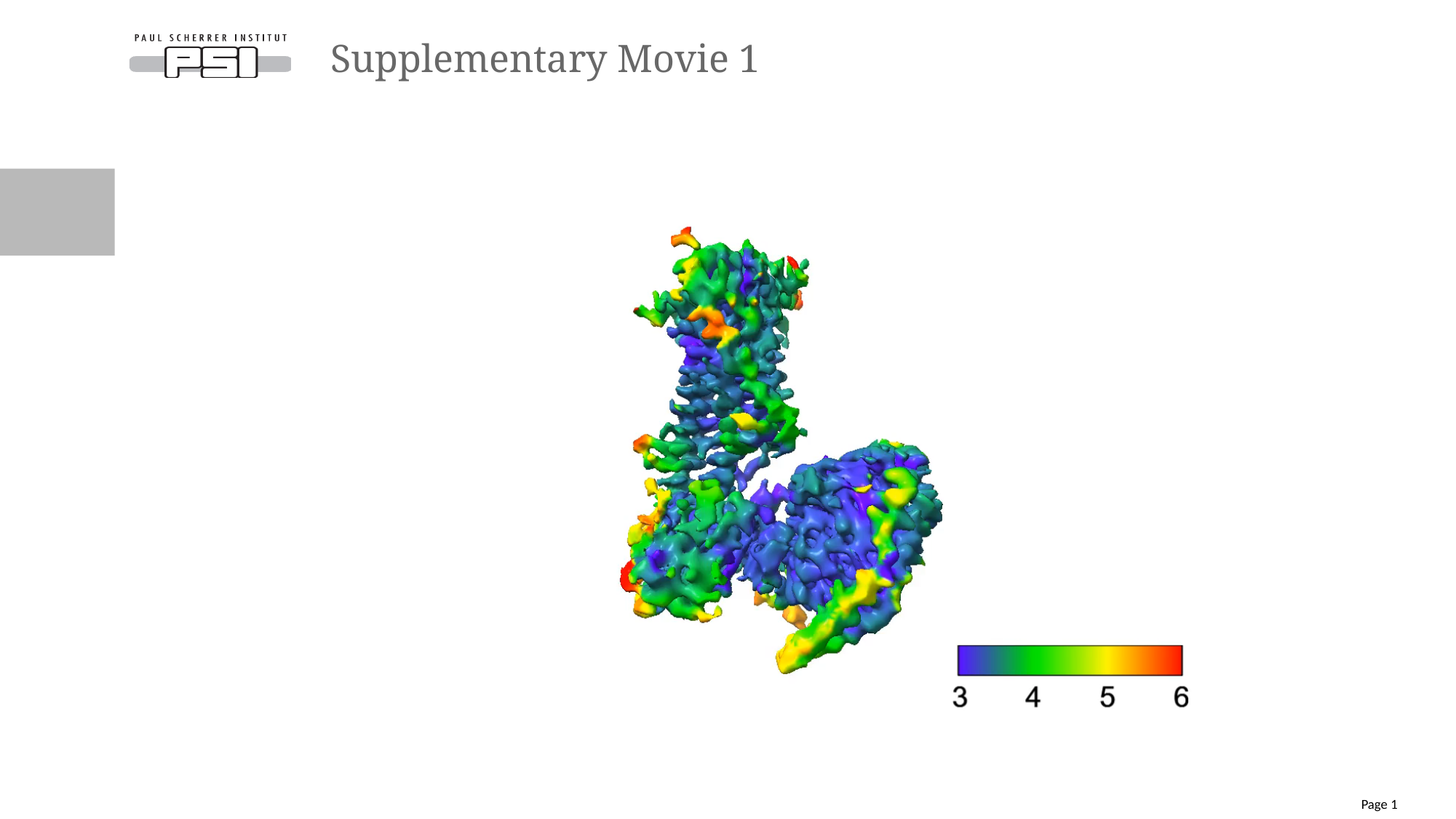

#### Supplementary Movie 1
Page 1

### Slide 2
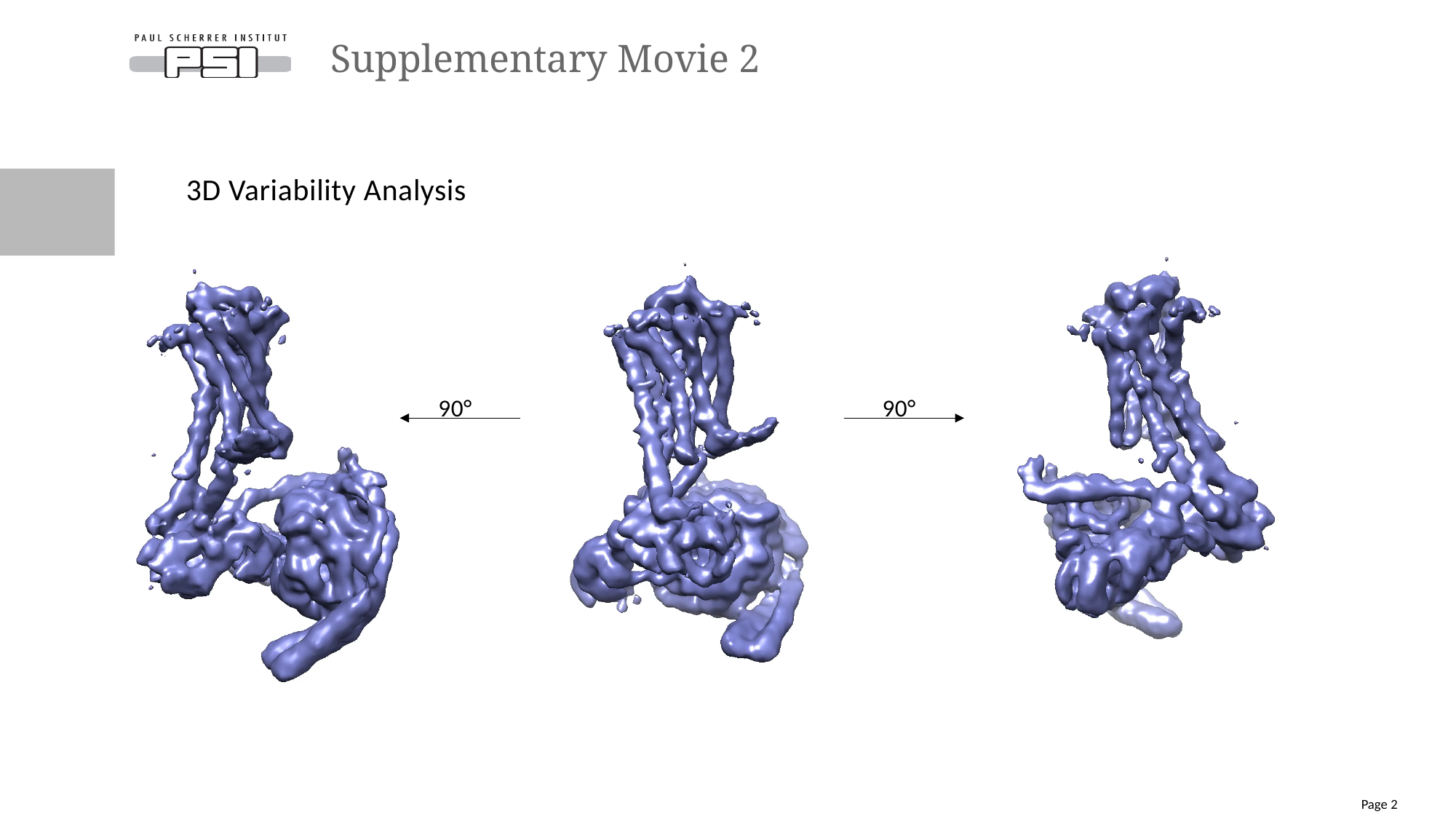

#### Supplementary Movie 2
3D Variability Analysis
90°
90°
Page 2

### Slide 3
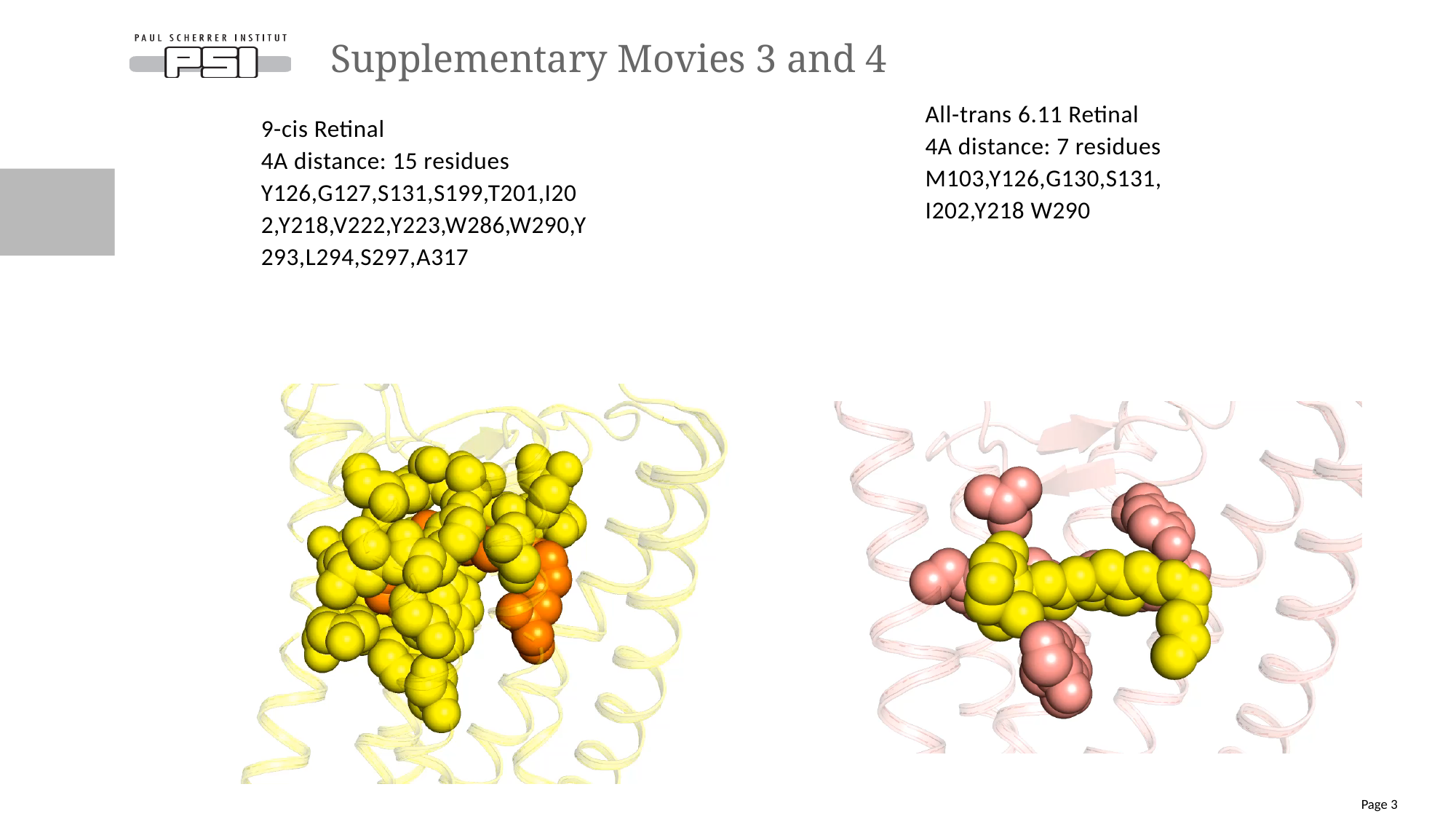

#### Supplementary Movies 3 and 4
All-trans 6.11 Retinal
4A distance: 7 residues
M103,Y126,G130,S131, I202,Y218 W290
9-cis Retinal
4A distance: 15 residues
Y126,G127,S131,S199,T201,I202,Y218,V222,Y223,W286,W290,Y293,L294,S297,A317
Page 3

### Slide 4
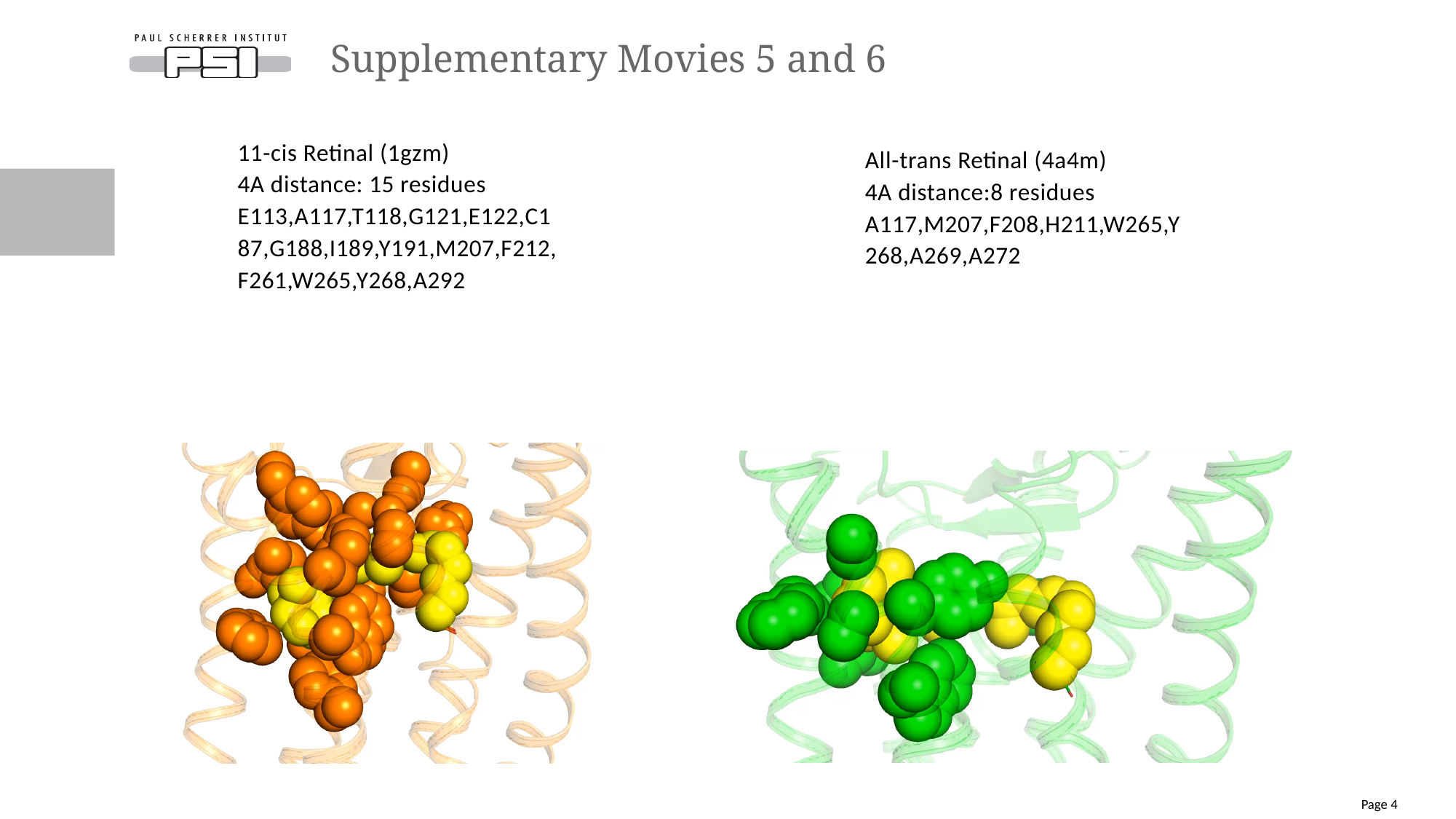

#### Supplementary Movies 5 and 6
11-cis Retinal (1gzm)
4A distance: 15 residues
E113,A117,T118,G121,E122,C187,G188,I189,Y191,M207,F212,F261,W265,Y268,A292
All-trans Retinal (4a4m)
4A distance:8 residues
A117,M207,F208,H211,W265,Y268,A269,A272
Page 4

### Slide 5
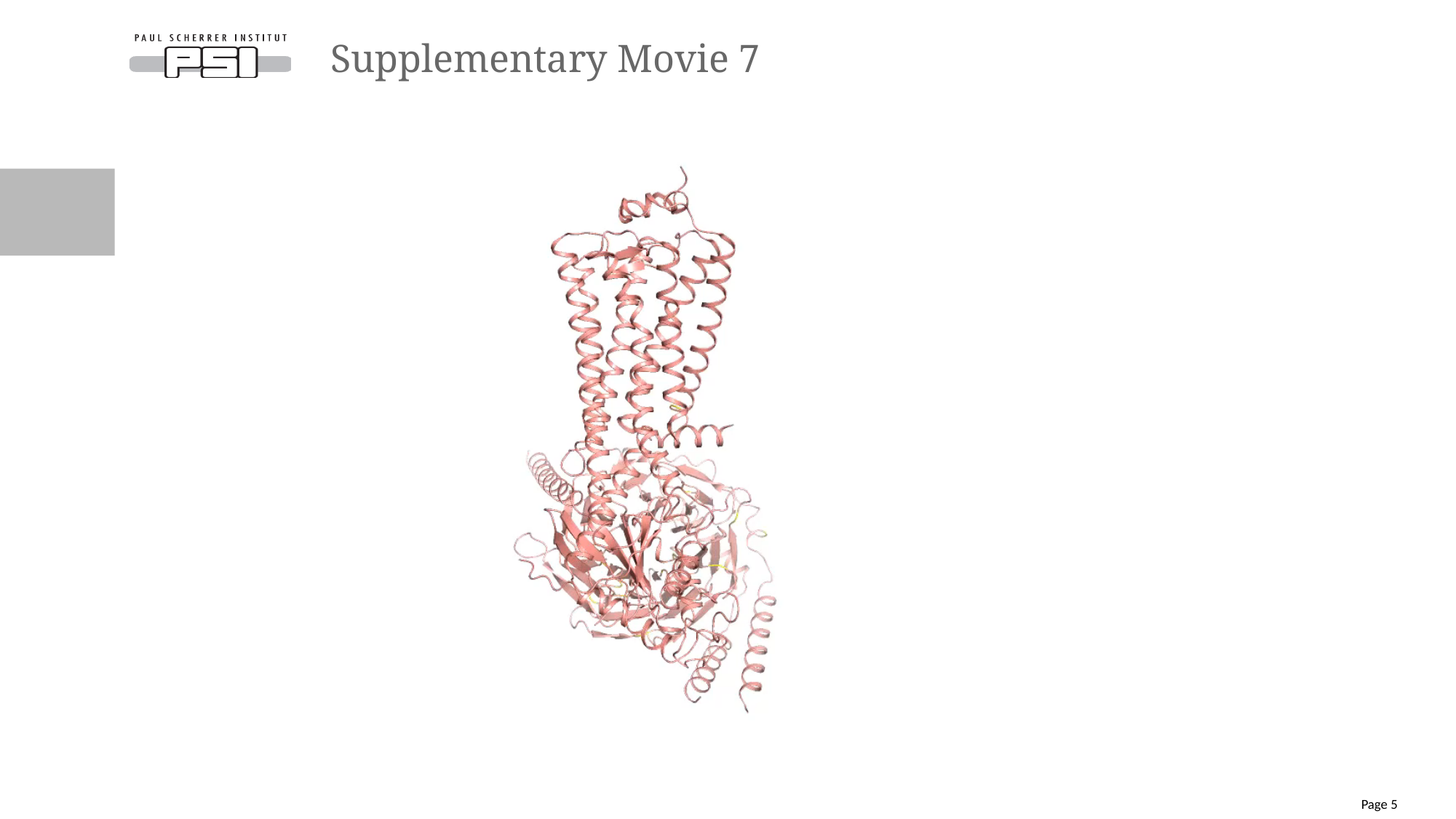

#### Supplementary Movie 7
Page 5

### Slide 6
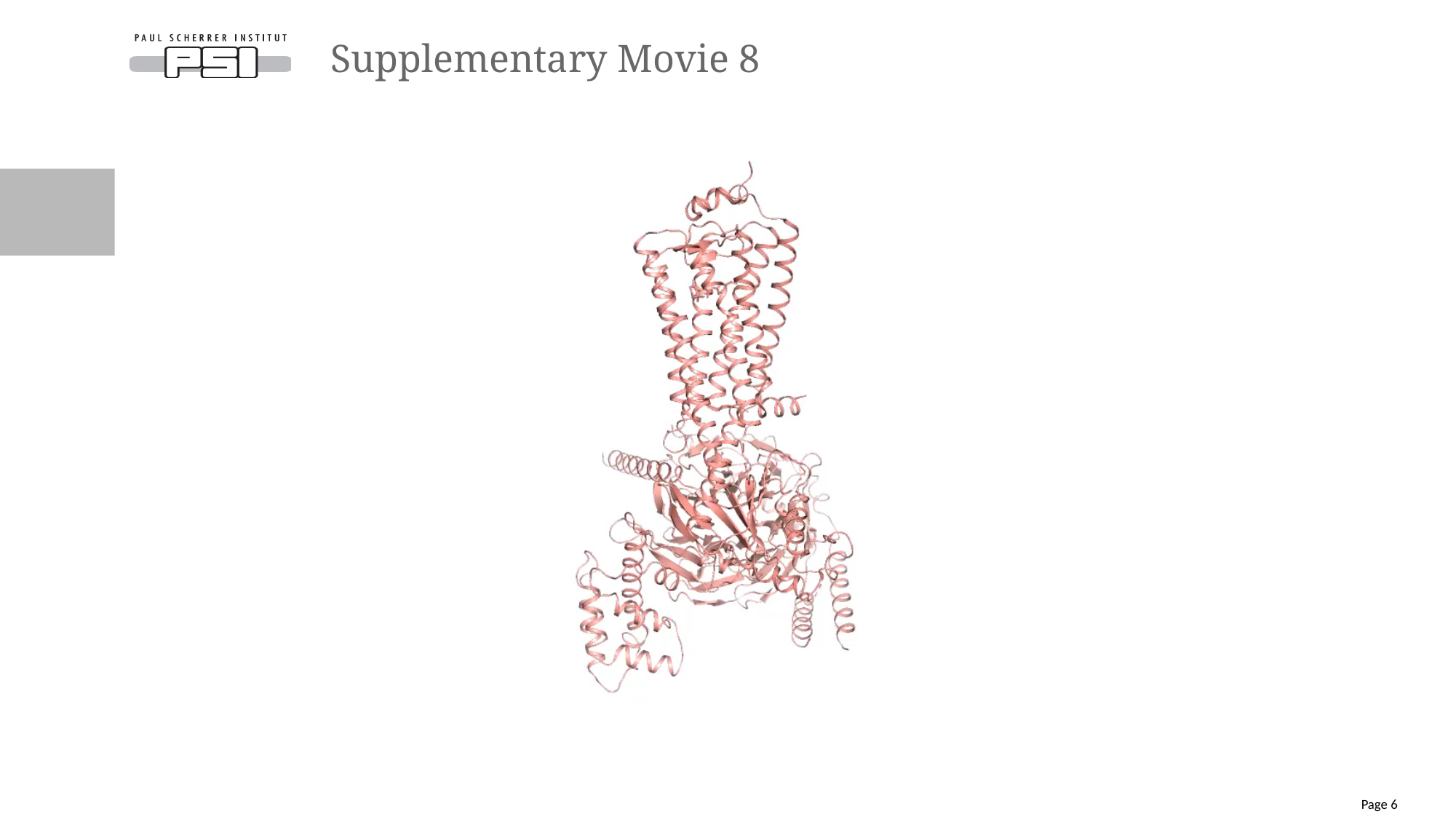

#### Supplementary Movie 8
Page 6
