## Supplementary Figures and Tables for "Active state structures of a bistable visual opsin bound to G proteins"

### Supplementary Figure 1

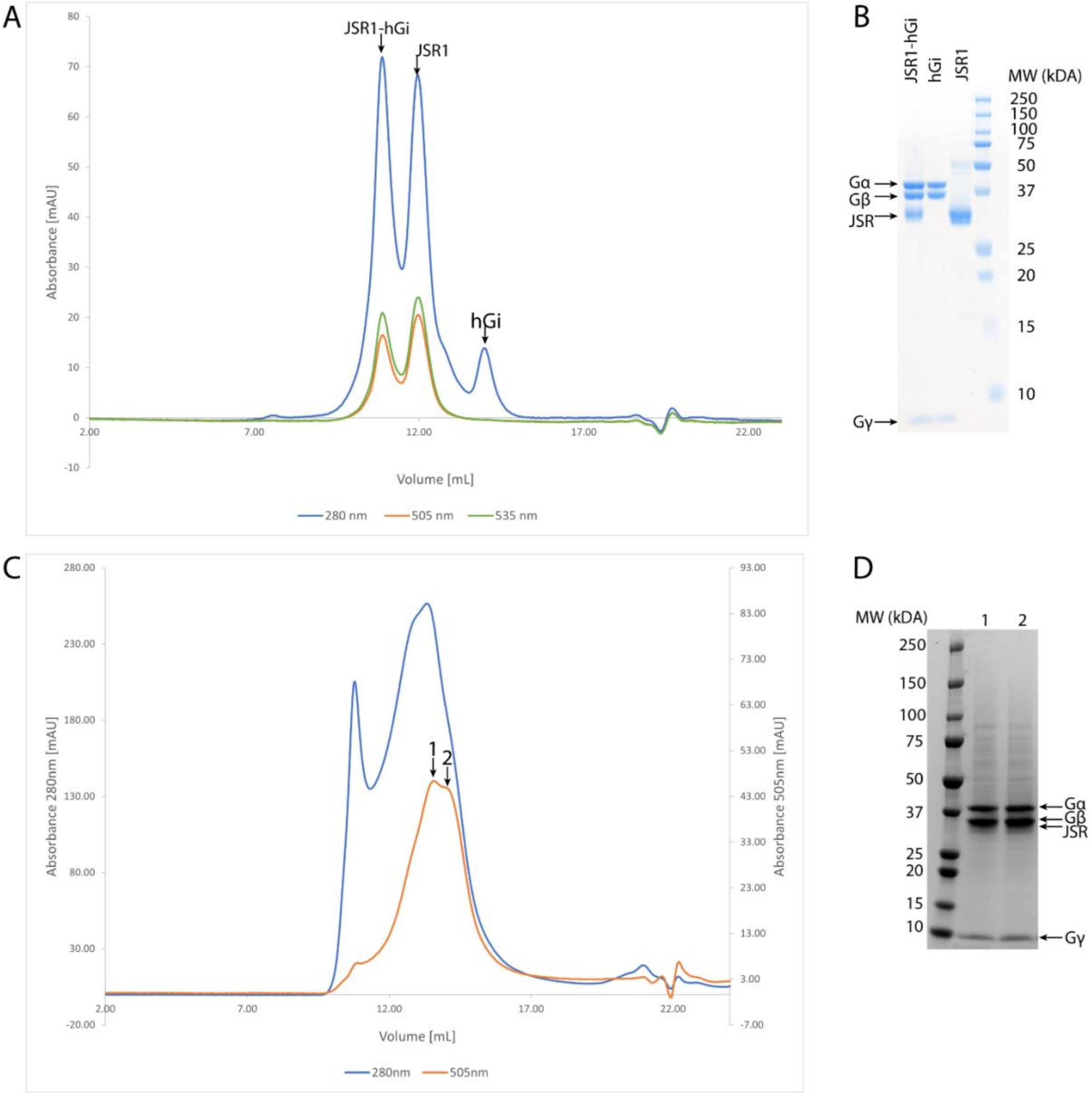


**Supplementary Figure 1: SEC and SDS-PAGE of the JSR1-hGi and JSR1-jsGiq complexes. A:** SEC chromatogram of the JSR1-hGi complex. Y-axis shows the absorbance in mAU and X-axis shows the volume in mL. **B:** Coomassie stained SDS-PAGE of the three peaks labeled in A. Individual subunits are labeled on the left side and the MW is indicated on the right side in kDa. **C:** SEC chromatogram of the JSR1-jsGiq complex. Y-axis shows the absorbance in mAU and X-axis shows the volume in mL. **D:** Coomassie stained SDS-PAGE of the two fractions indicated in C. Left side shows the MW in kDa and individual subunits are labeled on the right side. The bands for Gβ and JSR1 are overlapping.

**Supplementary Figure 2**

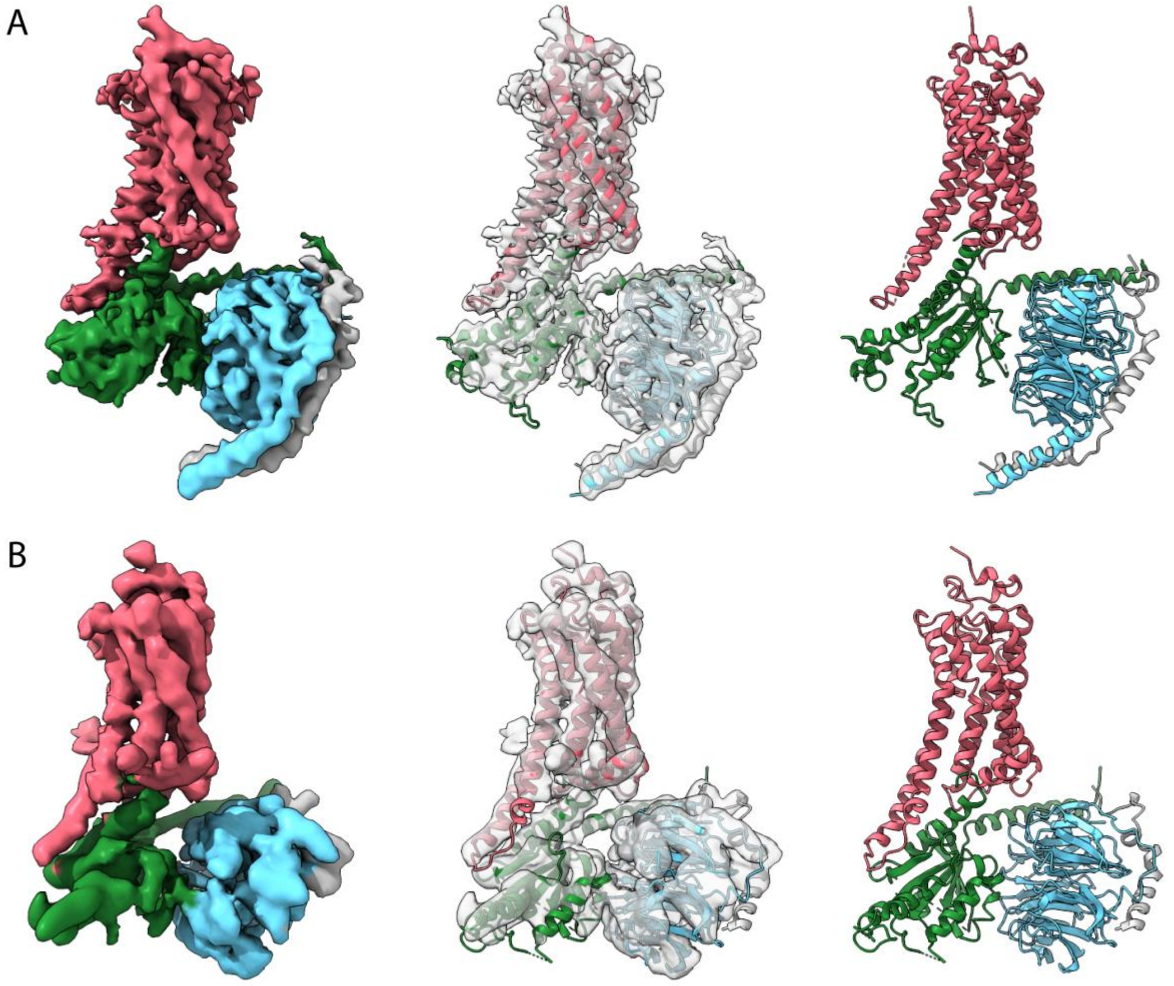


**Supplementary Figure 2: Cryo-EM maps and structural model for the JSR1-jsGiq_2 complex and the JSR1-hGi complex. A:** Left: Cryo-EM map colored by subunit. JSR1, jsGαiq, Gβ and Gγ subunits are shown in salmon, green, cyan and grey, respectively. Middle: Cryo-EM map is shown in grey and transparent with the model of JSR1-jsGiq_2 fitted into the map. Subunits of the model are colored in the same way as the map on the left. Right: Model of the JSR1-jsGiq_2 complex shown as cartoon. Subunits are colored as in the left cryo-EM map. **B:** Same figures as in A, colored in the same way, for the JSR1-hGi complex.

**Supplementary Figure 3**

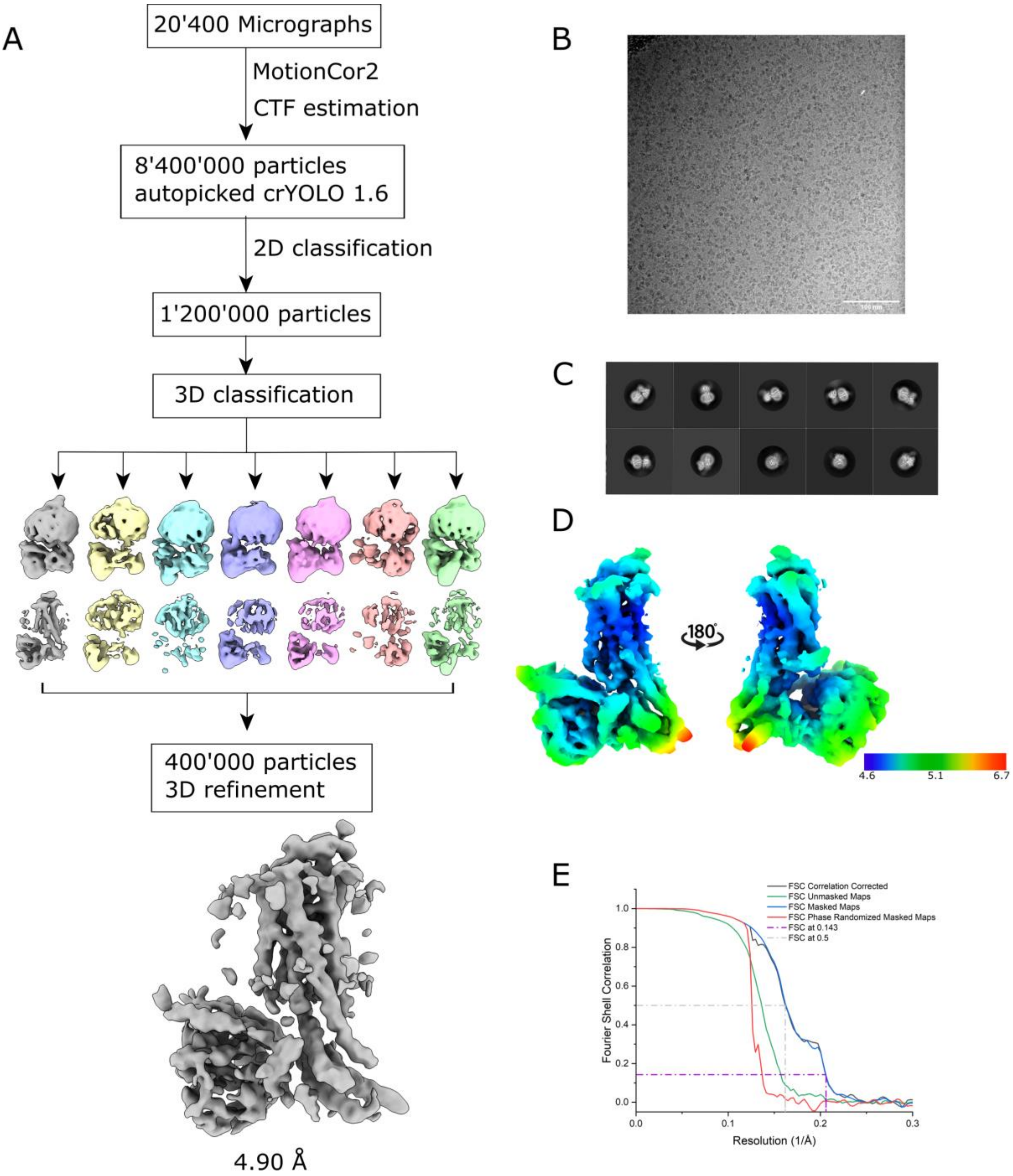


**Supplementary. Figure 3: Single-particle analysis of the JSR1-hGi complex. A**: Single-particle analysis workflow for the JSR1-hGi complex. **B:** Representative micrograph. **C:** 2D class averages for the JSR1-hGi complex in different orientations. **D:** 4.9 Å JSR1-hGi complex electron potential map colored by local resolution values. **E:** FSC plot for the JSR1-hGi complex at 4.9 Å resolution.

**Supplementary Figure 4**

A

| | H5.01 | H5.02 | H5.03 | H5.04 | H5.05 | H5.06 | H5.07 | H5.08 | H5.09 | H5.10 | H5.11 | H5.12 | H5.13 | H5.14 | H5.15 | H5.16 | H5.17 | H5.18 | H5.19 | H5.20 | H5.21 | H5.22 | H5.23 | H5.24 | H5.25 | H5.26 |
|---|---|---|---|---|---|---|---|---|---|---|---|---|---|---|---|---|---|---|---|---|---|---|---|---|---|---|
| Human Gi | T | K | N | V | Q | F | V | F | D | A | V | T | D | V | I | I | K | N | N | L | K | D | C | G | L | F |
| Human Gq | T | E | N | I | R | F | V | F | A | A | V | K | D | T | I | L | Q | L | N | L | K | E | Y | N | L | V |
| Hasarius Gq1 visual | T | E | N | I | R | L | V | F | C | A | V | K | D | T | I | L | Q | N | N | L | K | E | C | N | L | V |
| Hasarius Gq2 | T | E | N | I | R | F | V | F | A | A | V | K | D | T | I | L | Q | L | N | L | K | E | Y | N | L | V |

B

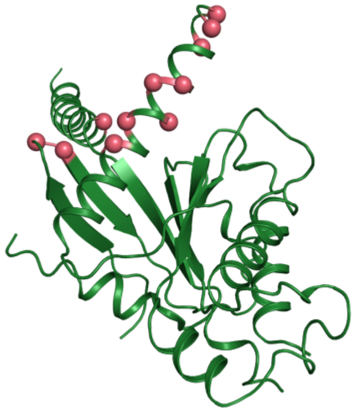

C

| Residue number (hGi) | Mutation |
|---|---|
| 31 | A→R |
| 193 | D→S |
| 194 | L→I |
| 337 | D→C |
| 340 | T→K |
| 342 | V→T |
| 344 | I→L |
| 345 | K→Q |
| 350 | D→E |
| 352 | G→N |
| 354 | F→V |

**Supplementary Figure 4: Sequence comparison and mutations introduced. A:** Table comparing the α5 helix sequences of the human Gαi, human Gαq, jumping spider Gαq1 visual (HaGq1: acc. No. LC799818) and jumping spider Gαq2 (HaGq2: acc. No. LC799819). **B:** Human Gαi structure from our JSR1-jsGiq complex. Locations where mutations were introduced are marked in red. **C:** Table with the mutations introduced to make the jsGiq chimera.

**Supplementary Figure 5**

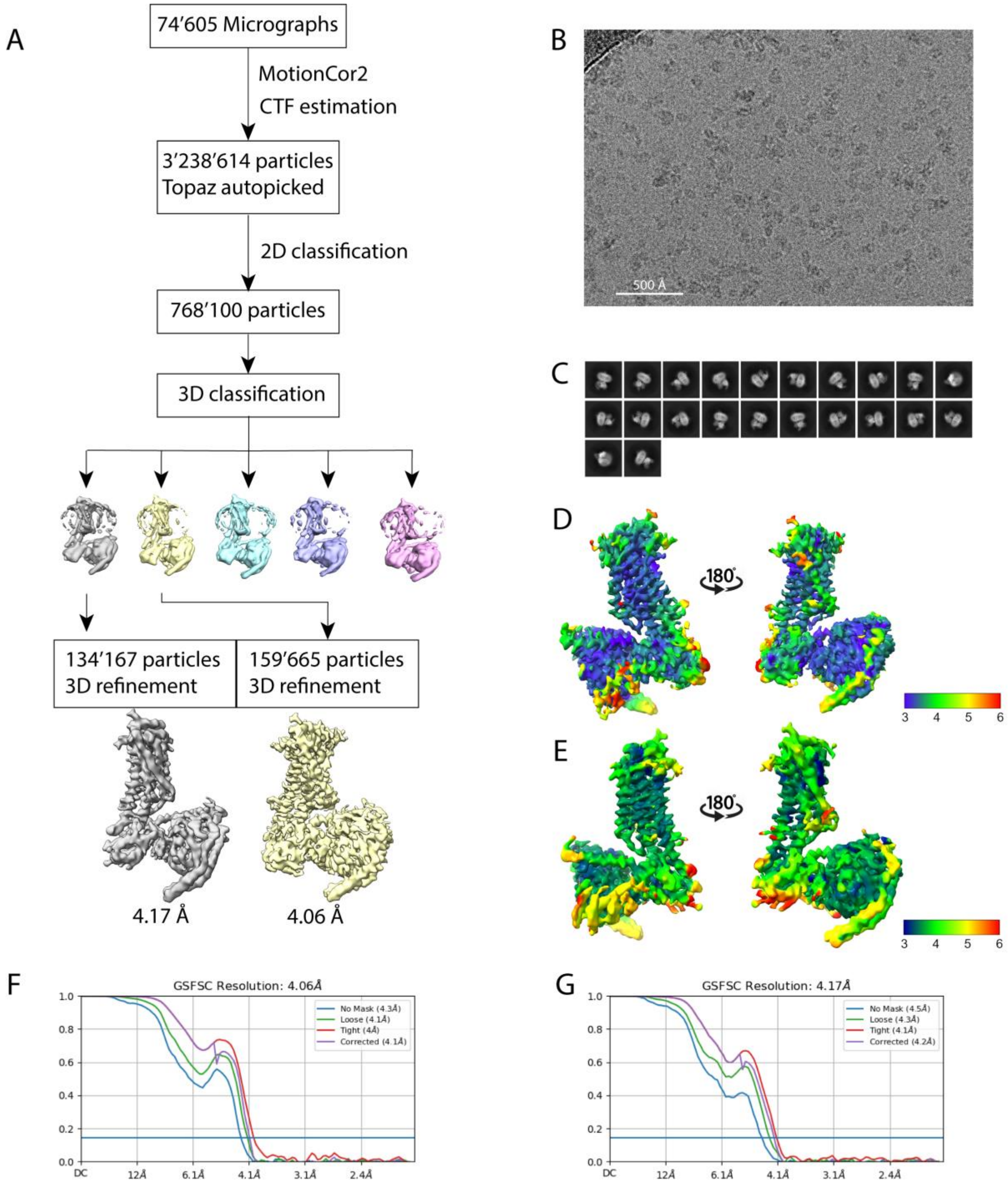


**Supplementary Figure 5: Single-particle analysis of the JSR1-jsGiq complex. A**: Single-particle analysis workflow for the JSR1-jsGiq complex. **B:** Representative micrograph taken at a nominal magnification of 165’000 (0.51 Å/pixel). **C:** Selected 2D class averages for the JSR1-jsGiq complex. **D:** 4.1 Å JSR1-jsGiq_1 complex electron potential map colored by local resolution values (Local Resolution Estimation, cryoSPARC). **E:** 4.2 Å JSR1-jsGiq_2 complex electron potential map colored by local resolution values (Local Resolution Estimation, cryoSPARC). **F:** FSC plot for the JSR1-jsGiq complex at 4.1 Å resolution. **G:** FSC plot for the JSR-jsGiq complex at 4.2 Å resolution.

**Supplementary Figure 6**

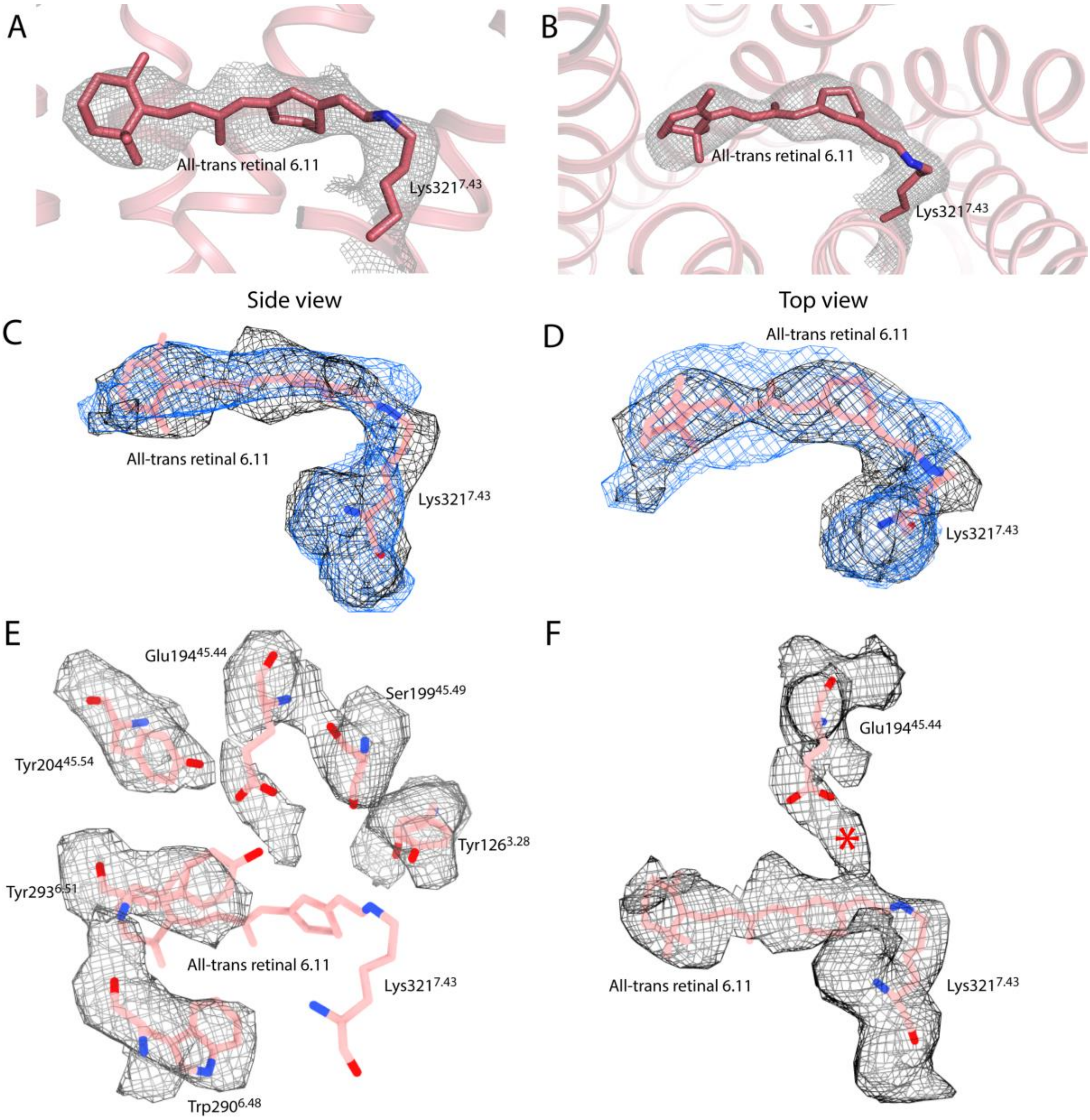


**Supplementary Figure 6: All-trans retinal 6.11 with the experimental cryo-EM map and comparison to the all-trans retinal from the JSR1-hGi complex. AB:** Visualization of the all-trans retinal 6.11 from the JSR1-jsGiq complex including the experimental cryo-EM map for the Lys321 and all-trans retinal 6.11 showed as an isomesh. A shows a side view and B shows a top view from the extracellular side. **CD:** Overlay of the all-trans retinal 6.11 with the electron density map from the JSR1-jsGiq complex (black mesh) and the electron density map of all-trans retinal from the JSR1-hGi complex (blue mesh). C shows a side view and D shows a top view from the extracellular side. **E:** Residues in the retinal environment shown with the experimental cryo-EM map overlaid as a mesh. **F:** Counterion (Glu194) and retinal with the Schiff base link with the experimental cryo-EM map overlaid. The red asterisk marks a potential water molecule position.

**Supplementary Figure 7**

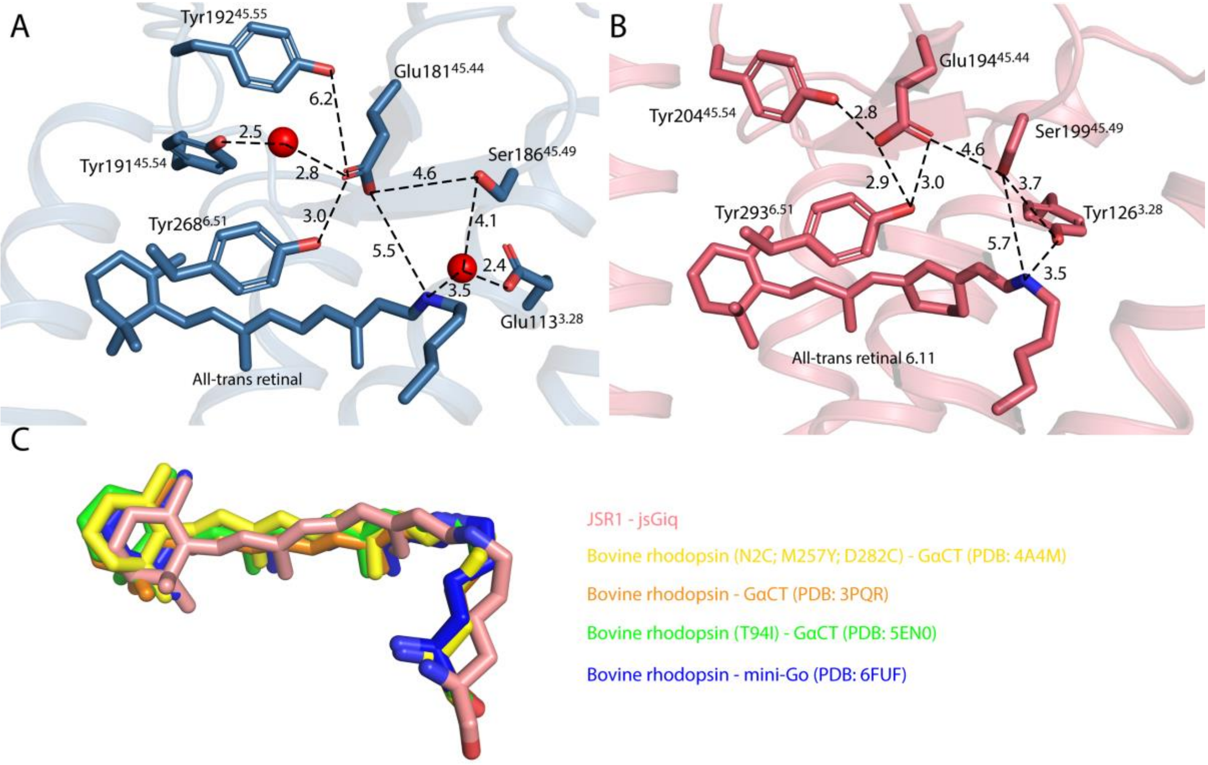


**Supplementary Figure 7: Retinal binding site comparison of bovine metarhodopsin-II and active state JSR1. A:** Retinal environment of bovine metarhodopsin II (PDB:5EN0). Important residues are shown as sticks. Carbon atoms are colored green, nitrogen atoms blue and oxygen atoms red. **B:** All-trans retinal 6.11 environment of the active JSR1. Corresponding residues from A are shown as sticks for JSR1 and carbon atoms are shown in salmon, nitrogen atoms in blue and oxygen atoms in red. **C:** Overlay of the all-trans retinal 6.11 from our JSR1 structure and all-trans retinal from four bovine metarhodopsin II structures (PDB: 4A4M (yellow); 3PQR (orange); 6FUF (blue); 5EN0 (green)).

**Supplementary Figure 8**

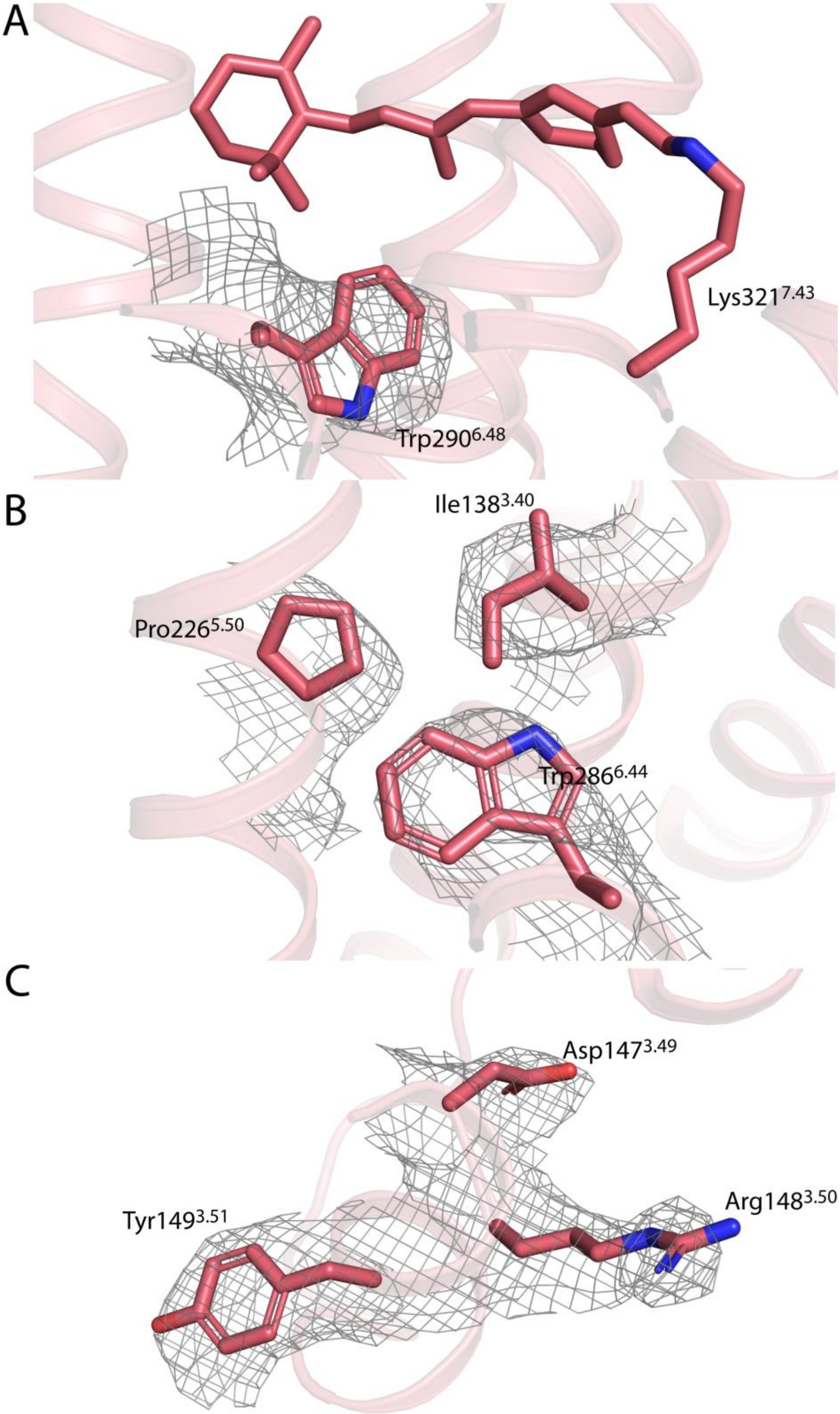


**Supplementary Figure 8: Microswitches with experimental cryo-EM map. A:** Trp290 from the C-W-x-P motif shown in stick representation with the experimental map shown as mesh. **B:** P-I-F motif with side chains shown as sticks and the experimental map shown as mesh. **C:** D-R-Y motif with side chains shown as sticks and the experimental map shown as mesh.

**Supplementary Figure 9**

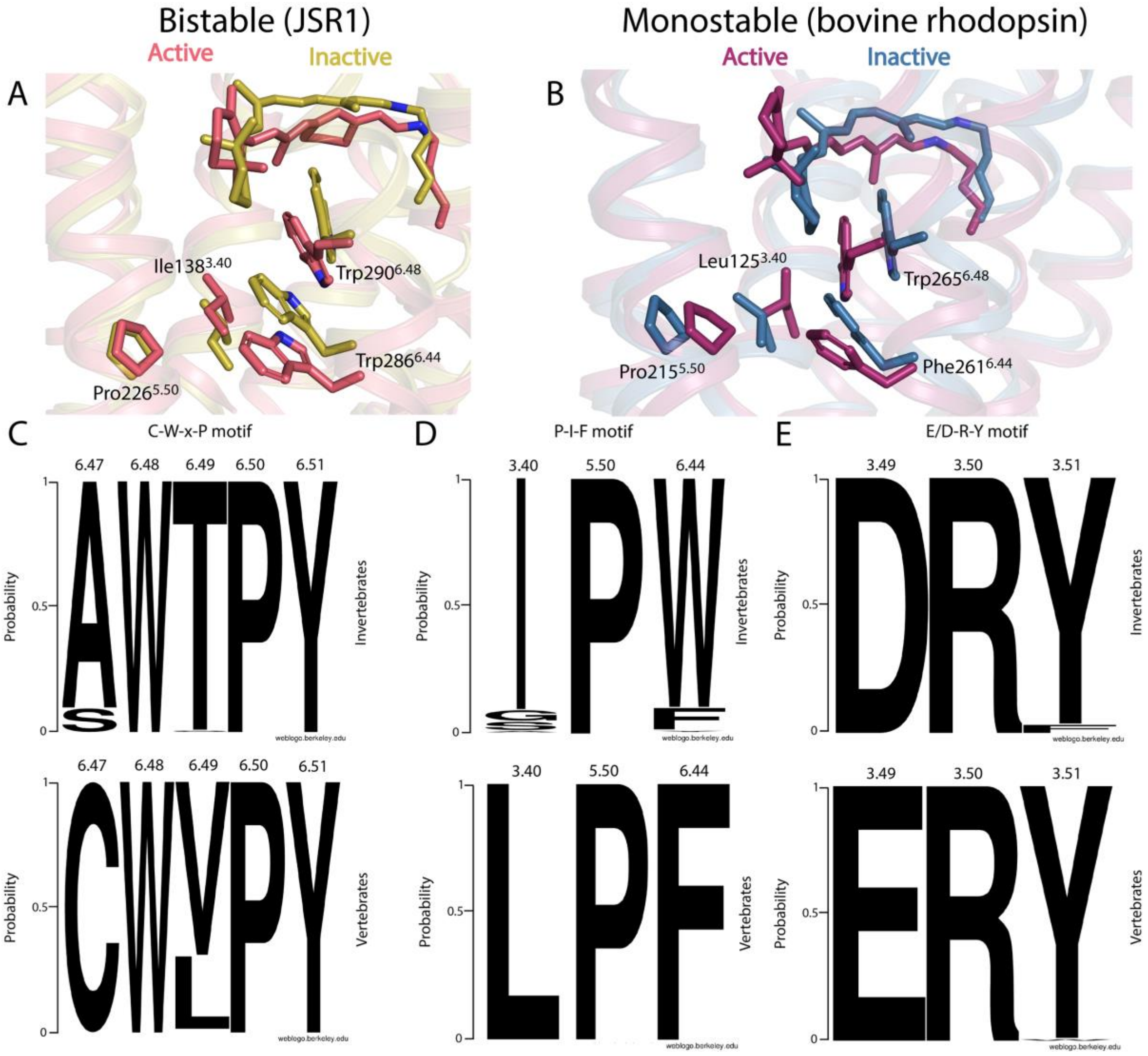


**Supplementary Figure 9: Activation of JSR1 compared to bovine rhodopsin with sequence conservation of microswitches in all opsins. A:** Conformational changes from the retinal isomerization to the P-I-F motif, leading to the outward movement of TM6. Inactive state JSR1 is shown is yellow (PDB: 6I9K) and active state JSR1 is shown in salmon. **B:** Conformational changes from the retinal isomerization to the P-I-F motif, leading to the outward movement of TM6. Inactive state bovine rhodopsin (PDB:1GZM) is shown in blue and active state bovine metarhodopsin II (PDB:5EN0) is shown in purple. **C:** Sequence conservation of the C-W-x-P motif in invertebrate rhodopsins (top) and vertebrate rhodopsins (bottom). **D:** Sequence conservation of the P-I-F motif in invertebrate rhodospins (top) and vertebrate rhodospins (bottom). **E:** Sequence conservation of the E/D-R-Y motif in invertebrate rhodospins (top) and vertebrate rhodopsins (bottom).

**Supplementary Figure 10**

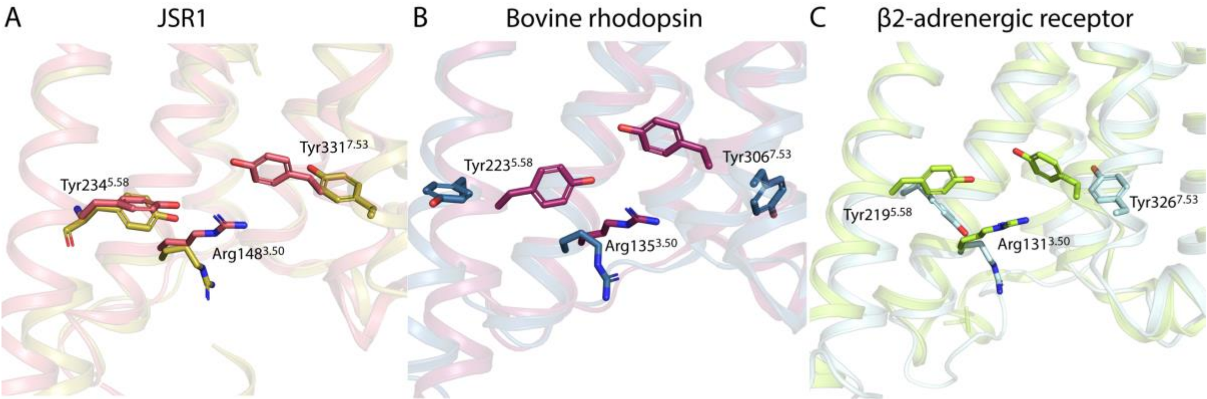


**Supplementary Figure 10: Conformational changes of Arg$^{3.50}$, Tyr$^{5.58}$ and Tyr$^{7.53}$ upon receptor activation. A:** Inactive JSR1 is shown in yellow and active JSR1 in salmon. Arg148$^{3.50}$, Tyr234$^{5.58}$ and Tyr331$^{7.53}$ are shown as sticks in the respective colour. Oxygen atoms are shown in red and nitrogen atoms in blue. **B:** Inactive bovine rhodopsin is shown in blue and active bovine rhodopsin in purple. Arg135$^{3.50}$, Tyr223$^{5.58}$ and Tyr306$^{7.53}$ are shown as sticks in the respective colour. Oxygen atoms are shown in red and nitrogen atoms in blue. **C:** Inactive β2-adrenergic receptor is shown in palecyan and active β2-adrenergic receptor is shown in palegreen. Arg131$^{3.50}$, Tyr219$^{5.58}$ and Tyr326$^{7.53}$ are shown as sticks in the respective colour. Oxygen atoms are shown in red and nitrogen atoms in blue.

**Supplementary Figure 11**

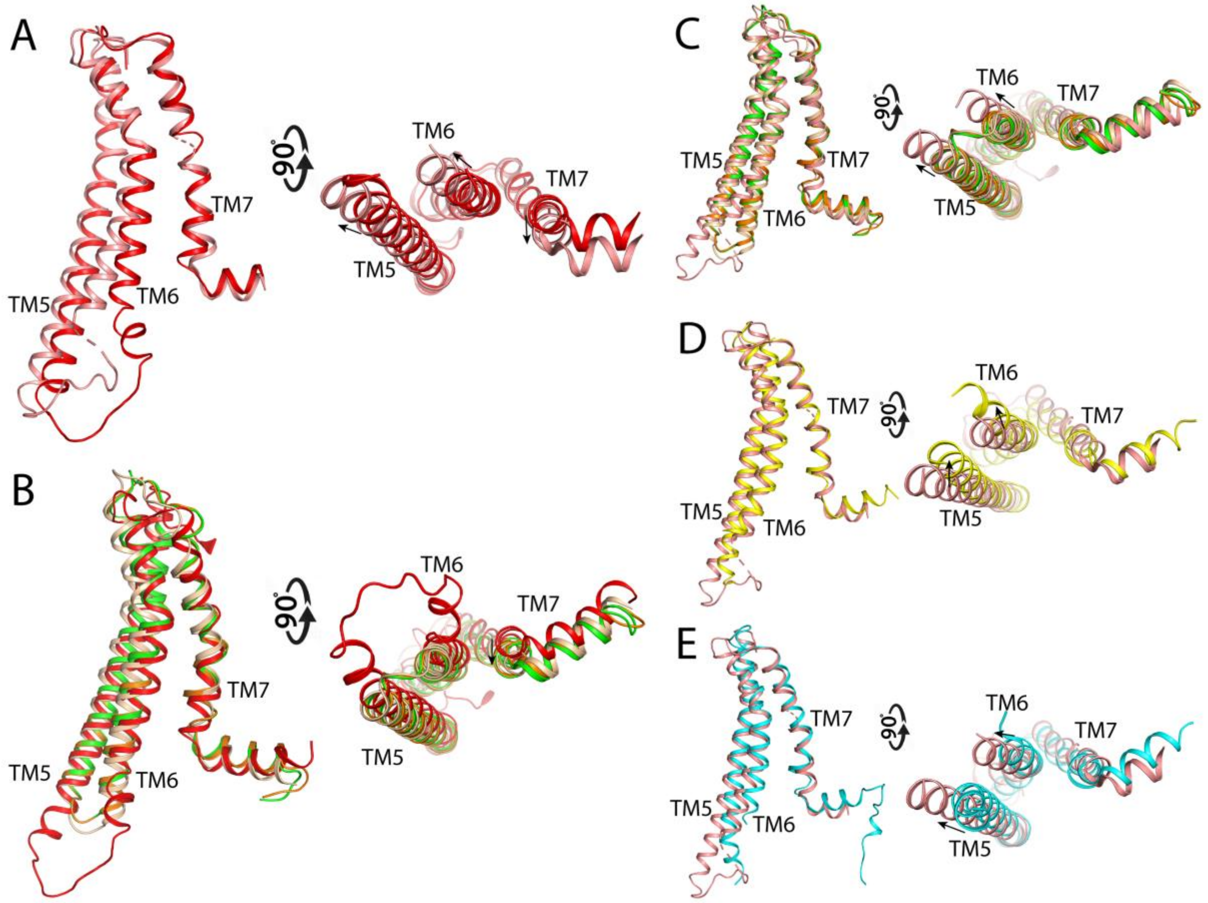


**Supplementary Figure 11: TM5/6 and TM7 relocation comparison. A:** Overlay of TM5/6/7 of the JSR1-jsGiq_1 complex (salmon) and the JSR1-hGi complex (red). **B:** Overlay of TM5/6/7 of the JSR1-hGi complex with bovine metarhodopsin II bound to GαCT (PDB: 3PQR (orange); PDB: 4A4M (green)) and the rhodopsin-hGi complex (PDB:6CMO (wheat)). **C:** Overlay of TM5/6/7 of the JSR1-jsGiq_1 complex with bovine metarhodopsin II bound to GαCT (PDB: 3PQR (orange); PDB: 4A4M (green)) and the rhodopsin-hGi structure (PDB:6CMO (wheat)). **D:** Overlay of TM5/6/7 of the JSR1-jsGiq_1 complex (salmon) with the β2-adrenergic receptor-Gs complex (PDB: 3SN6, yellow). **E:** Overlay of TM5/6/7 of the JSR1-jsGiq_1 complex (salmon) with the M1 receptor-G11 complex (PDB: 6OIJ, cyan).

**Supplementary Figure 12**

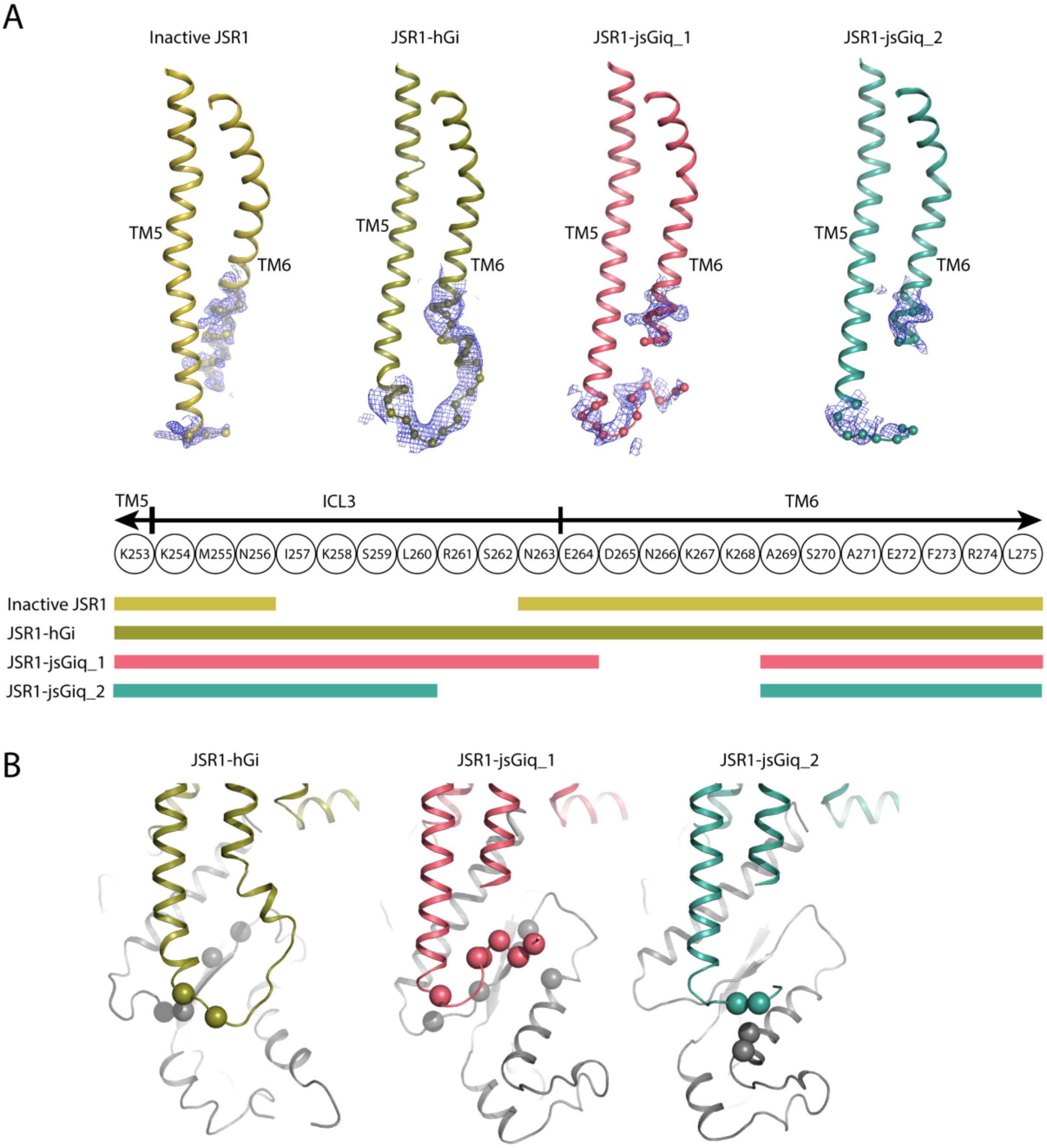


**Supplementary Figure 12: Comparison of TM5-ICL3-TM6 in the JSR1 structures. A:** The top panel shows structures of TM5-ICL3-TM6 of inactive JSR1 (PDB=6i9k), active JSR1 from the JSR1-hGi model, active JSR1 from the JSR1-jsGiq_1 model and active JSR1 from the JSR1-jsGiq_2 model. The experimental cryo-EM maps are shown for residues 253-275 of JSR1. The bottom panel shows a schematic representation of the residues 253-275. The colored rows show which residues are present (solid bar) and missing (no bar) from the respective structures**. B:** Observed contact between JSR1 ICL3 and the α4-β6 region of Gα. Based on the built structural models, residues of JSR1 ICL3 within 4 Å to the α4-β6 region of Gα (and vice versa) are plotted in sphere style for their Cα. JSR1 is colored in olive (JSR1-hGi), salmon (JSR1-jsGiq_1) and teal (JSR1-jsGiq_2), respectively. Gα is colored in grey.

**Supplementary Figure 13**

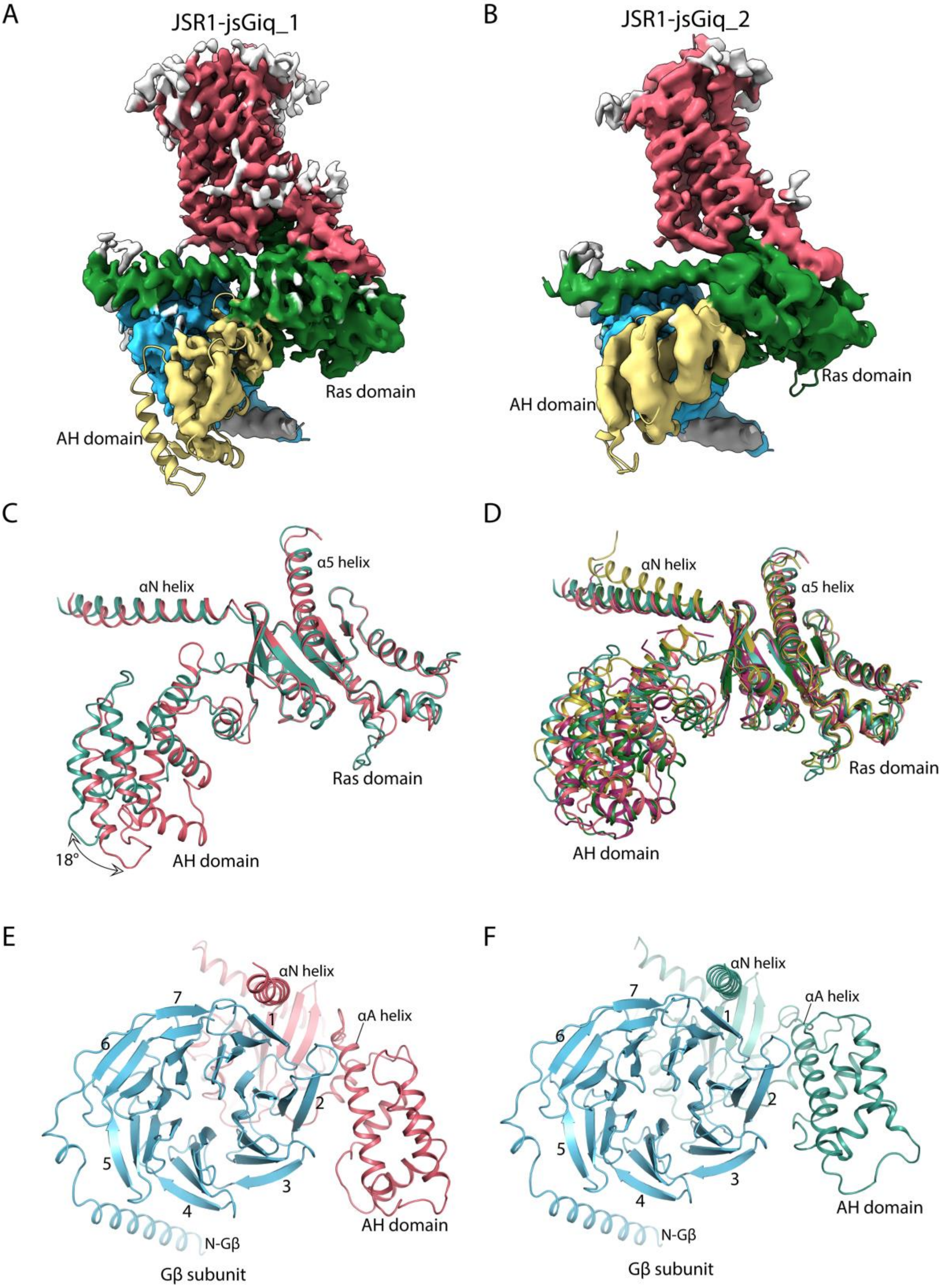


**Supplementary Figure 13: A, B:** Experimental cryo-EM maps of the JSR1-jsGiq complex structures 1 (A) and 2 (B) overlaid with their models. Specific colors are assigned for JSR1 (salmon), Gα Ras domain (green), Gα AH domain (yellow), Gβ (cyan) and Gγ (grey). **C:** Structure comparison of the jsGαiq subunit from the JSR1-jsGiq_1 (salmon) and JSR1-jsGiq_2 (teal) complexes. The α-helical domain (AH domain), αN helix, α5 helix and the Ras domain are labelled. **D:** Structure comparison of the Gα subunit from the JSR1-jsGiq_1 (salmon), JSR1-jsGiq_2 (teal), GABA(B) receptor-hGi (PDB: 7EB2; yellow), NTSR1-hGi (PDB: 7L0S; purple) and the cannabinoid receptor 2-hGi (PDB: 6PT0; green) complex structures. The α-helical domain (AH domain), αN helix, α5 helix and the Ras domain are labelled. **E:** Structure of the jsGαiq and Gβ subunits from the JSR1-jsGiq_1 complex. The Gα subunit is colored in salmon and the Gβ subunit in cyan. The AH domain, αN helix and the H.HA helix of the jsGαiq are labelled. The numbers of the Gβ subunit represent the numbering of the β blades according to Wall *et al.* [1]. **F:** Structure of the jsGαiq and Gβ subunits from the JSR1-jsGiq_2 complex. The Gα subunit is colored in teal and the Gβ subunit in cyan. The AH domain, αN helix and the H.HA helix of the jsGαiq are labelled. The numbers of the Gβ subunit represent the numbering of the β blades according to Wall *et al.* [1].

**Supplementary Figure 14**

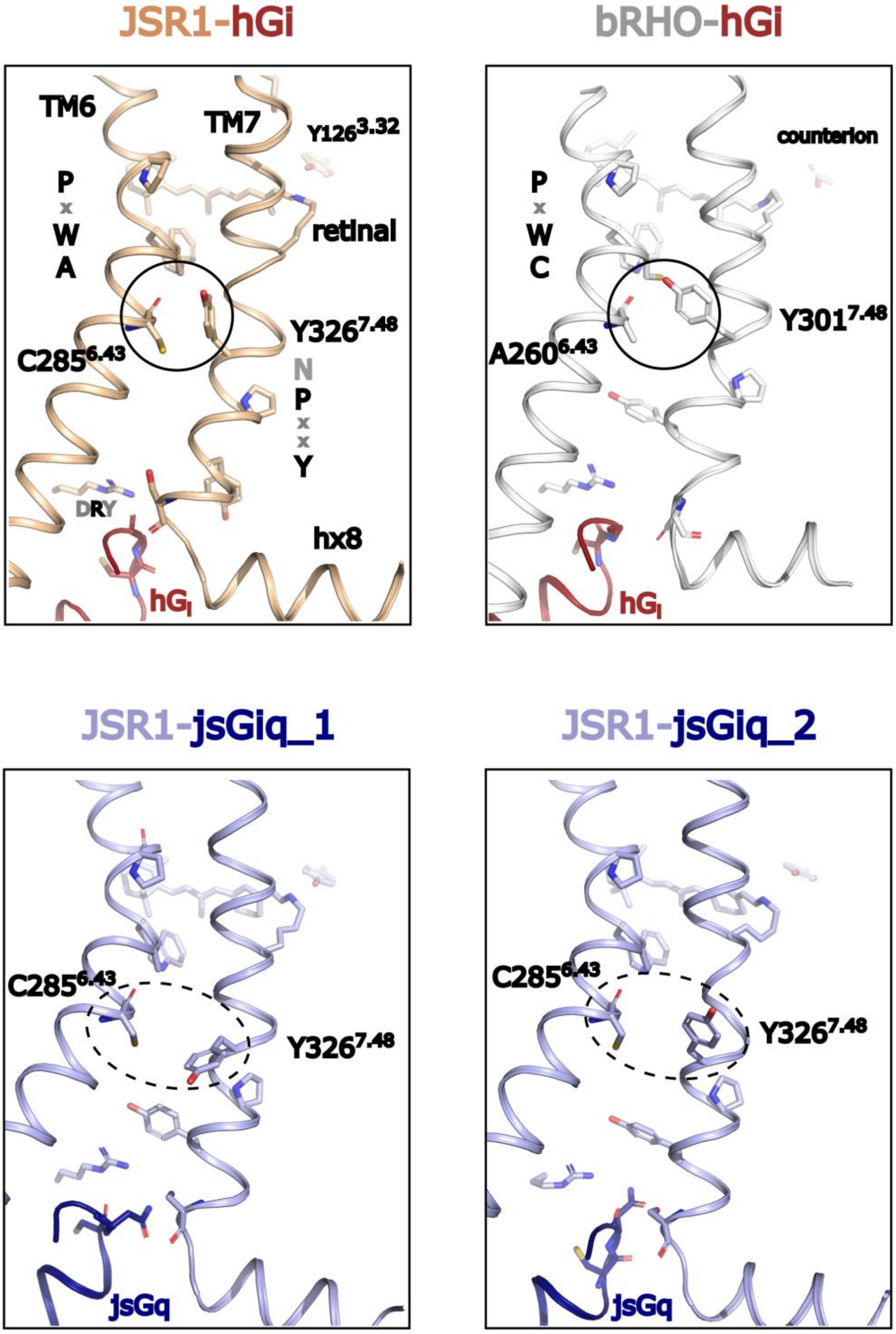


**Supplementary Figure 14:** Detail of TM6/TM7/H8 in the JSR1-hGi complex (top left), the two conformations of the JSR1-jsGiq complex (bottom), and the bovine rhodopsin-hGi complex (top right). Top panels: The packing at the 6.43/7.48 residues (circle) is located right between the A/C-W-x-P motif in TM6 and the N-P-x-x-Y motif in TM7, and is similar in JSR1 and rhodopsin bound to hGi. In bovine rhodopsin, the packing is stronger, including a inter-helical hydrogen bond. Bottom panels: This packing is looser in jsGiq-bound JSR1.

1. Wall, M.A., et al., *The structure of the G protein heterotrimer Gi alpha 1 beta 1 gamma 2.* Cell, 1995. **83**(6): p. 1047-58.

| **Residue** | R148 (3.50) | V151 (3.53) | I152 (3.54) | M156 (34.51) | A157 (34.52) | P160 (34.55) | I237 (5.61) | H244 (5.68) | L248 (5.72) | Q251 (5.75) | K254 (5.78) | M255 (5.79) | L260 (ICL3) | R261 (ICL3) | S262 (ICL3) | N263 (ICL3) | R274 (6.32) | L275 (6.33) | V278 (6.36) | S334 (7.56) | H335 (8.47) | P336 (8.48) | K337 (8.49) |
|---|---|---|---|---|---|---|---|---|---|---|---|---|---|---|---|---|---|---|---|---|---|---|---|
| R31 (G.hns1.02) | | | | orange | green | | | | | | | | | | | | | | | | | | |
| E308 (G.H4.16) | | | | | | | | | | | | | | | green | | | | | | | | |
| E318 (G.h4s6.12) | | | | | | | | | | | | | | orange | | green | | | | | | | |
| I319 (G.h4s6.13) | | | | | | | | | | | | | | | green | | | | | | | | |
| H322 (G.S6.04) | | | | | | | | | | | | green | | | | | | | | | | | |
| C325 (G.s6h5.02) | | | | | | | | | | | | green | | | | | | | | | | | |
| Q333 (G.H5.5) | | | | | | | | | | orange | green | | | | | | | | | | | | |
| F334 (G.H5.6) | | | | | | | | | | | | | green | | | | | | | | | | |
| C337 (G.H5.9) | | | | | | | | | | green | | | | | | | | | | | | | |
| K340 (G.H5.12) | | | | | | | | green | | | | | | | | | | | | | | | |
| D341 (G.H5.13) | | | | | | | | green | | | | | | orange | | | | | | | | | |
| N347 (G.H5.19) | | | green | | | | | | | | | | | | | | | | | | | | |
| L348 (G.H5.20) | | | green | | | | | | | | | | | | | | | | | | | | |
| K349 (G.H5.21) | | | | | | | | | | | | | | | | | | | | | | green | |
| E350 (G.H5.22) | | | | | | | | | | | | | | | | | | | | | | | orange |
| C351 (G.H5.23) | orange | green | | | | | | | | | | | | | | | | | | | | | |
| N352 (G.H5.24) | green | | | | | | | | | | | | | | | | | | | orange | orange | green | |
| L353 (G.H5.25) | green | | | | | | green | | | | | | | | | | | green | green | | | | |
| V354 (G.H5.26) | | | | | | | | | | | | | | | | | orange | | green | | | | |

**Supplementary Table 1:** JSR1-jsGiq_1 interaction table. Top row shows the JSR1 residue number and GPCR numbering. Left column shows the jsGiq residue number and G protein numbering. Orange fields symbolize polar interactions and green fields residues within 4 Å distance.

| Residue | R148 (3.50) | V151 (3.53) | I152 (3.54) | M156 (34.51) | A157 (34.52) | P160 (34.55) | H244 (5.68) | L248 (5.72) | Q251 (5.75) | K254 (5.78) | M255 (5.79) | I257 (ICL3) | K258 (ICL3) | L260 (ICL3) | R274 (6.32) | V278 (6.36) | Y331 (7.53) | S334 (7.56) | H335 (8.47) | P336 (8.48) | K337 (8.49) |
|---|---|---|---|---|---|---|---|---|---|---|---|---|---|---|---|---|---|---|---|---|---|
| R31 (G.hns1.02) | | | | | | | | | | | | | | | | | | | | | |
| I194 (G.S3.01) | | | | | | | | | | | | | | | | | | | | | |
| E298 (G.H4.05) | | | | | | | | | | | | | | | | | | | | | |
| A301 (G.H4.08) | | | | | | | | | | | | | | | | | | | | | |
| C337 (G.H5.9) | | | | | | | | | | | | | | | | | | | | | |
| K340 (G.H5.12) | | | | | | | | | | | | | | | | | | | | | |
| D341 (G.H5.13) | | | | | | | | | | | | | | | | | | | | | |
| I343 (G.H5.15) | | | | | | | | | | | | | | | | | | | | | |
| K349 (G.H5.21) | | | | | | | | | | | | | | | | | | | | | |
| E350 (G.H5.22) | | | | | | | | | | | | | | | | | | | | | |
| N352 (G.H5.24) | | | | | | | | | | | | | | | | | | | | | |
| L353 (G.H5.25) | | | | | | | | | | | | | | | | | | | | | |
| V354 (G.H5.26) | | | | | | | | | | | | | | | | | | | | | |

**Supplementary Table 2:** JSR1-jsGiq_2 interaction table. Top row shows the JSR1 residue number and GPCR numbering. Left column shows the jsGiq residue number and G protein numbering. Orange fields symbolize polar interactions and green fields residues within 4 Å distance.

| Protein | Sequence | Sources |
|---|---|---|
| JSR1 | MLPHAAKMAARVAGDHDGRNISIVDLLPEDMLPMIHEHWYKFPPME<br>TSMHYILGMLIIVIGIISVSGNGVVMYLMMTVKNLRTPGNFLVLNL<br>ALSDFGMLFFMMPTMSINCFAETWVIGPFMCELYGMIGSLFGSASI<br>WSLVMITLDRYNVIVKGMAGKPLTKVGALLRMLFVWIWSLGWTIAP<br>MYGWSRYVPEGSMTSCTIDYIDTAINPMSYLIAYAIFVYFVPLFII<br>IYCYAFIVMQVAAHEKSLREQAKKMNIKSLRSNEDNKKASAEFRLA<br>KVAFMTICCWFMAWTPYLTLSFLGIFSDRTWLTPMTSVWGAIFAKA<br>SACYNPIVYGISHPKYRAALHDKFPCLKCGSDSPKGDSASTVAESE<br>KAGEETSQVAPA | Expression in HEK293 GnTI- |
| Human Gαi1 | MKKHHHHHHHHHHENLYFQGGSMGCTLSAEDKAAVERSKMIDRNLR<br>EDGEKAAREVKLLLLGAGESGKSTIVKQMKIIHEAGYSEEECKQYK<br>AVVYSNTIQSIIAIIRAMGRLKIDFGDSARADDARQLFVLAGAAEE<br>GFMTAELAGVIKRLWKDSGVQACFNRSREYQLNDSAAYYLNDLDRI<br>AQPNYIPTQQDVLRTRVKTTGIVETHFTFKDLHFKMFDVGGQRSER<br>KKWIHCFEGVTAIIFCVALSDYDLVLAEDEEMNRMHESMKLFDSIC<br>NNKWFTDTSIILFLNKKDLFEEKIKKSPLTICYPEYAGSNTYEEAA<br>AYIQCQFEDLNKRKDTKEIYTHFTCATDTKNVQFVFDAVTDVIIKN<br>NLKDCGLF | Expression in E. coli |
| jsGαiq | MGCTLSAEDKAAVERSKMIDRNLREDGEKARREVKLLLLGAGESGK<br>STIVKQMKIIHEAGYSEEECKQYKAVVYSNTIQSIIAIIRAMGRLK<br>IDFGDSARADDARQLFVLAGAAEEGFMTAELAGVIKRLWKDSGVQA<br>CFNRSREYQLNDSAAYYLNDLDRIAQPNYIPTQQDVLRTRVKTTGI<br>VETHFTFKSIHFKMFDVGGQRSERKKWIHCFEGVTAIIFCVALSDY<br>DLVLAEDEEMNRMHESMKLFDSICNNKWFTDTSIILFLNKKDLFEE<br>KIKKSPLTICYPEYAGSNTYEEAAAYIQCQFEDLNKRKDTKEIYTH<br>FTCATDTKNVQFVFCAVKDTILQNNLKECNLV | Expression in Hi5 |
| Bovine Gβ1 | MSELDQLRQEAEQLKNQIRDARKACADATLSQITNNIDPVGRIQMR<br>TRRTLRGHLAKIYAMHWGTDSRLLVSASQDGKLIIWDSYTTNKVHA<br>IPLRSSWVMTCAYAPSGNYVACGGLDNICSIYNLKTREGNVRVSRE<br>LAGHTGYLSCCRFLDDNQIVTSSGDTTCALWDIETGQQTTTFTGHT<br>GDVMSLSLAPDTRLFVSGACDASAKLWDVREGMCRQTFTGHESDIN<br>AICFFPNGNAFATGSDDATCRLFDLRADQELMTYSHDNIICGITSV<br>SFSKSGRLLLAGYDDFNCNVWDALKADRAGVLAGHDNRVSCLGVTD<br>DGMAVATGSWDSFLKIWN | Extraction from Bovine retina |
| Human Gβ1 | MHHHHHHHHHHLEVLFQGPGSSGSELDQLRQEAEQLKNQIRDARKA<br>CADATLSQITNNIDPVGRIQMRTRRTLRGHLAKIYAMHWGTDSRLL<br>VSASQDGKLIIWDSYTTNKVHAIPLRSSWVMTCAYAPSGNYVACGG<br>LDNICSIYNLKTREGNVRVSRELAGHTGYLSCCRFLDDNQIVTSSG<br>DTTCALWDIETGQQTTTFTGHTGDVMSLSLAPDTRLFVSGACDASA<br>KLWDVREGMCRQTFTGHESDINAICFFPNGNAFATGSDDATCRLFD<br>LRADQELMTYSHDNIICGITSVSFSKSGRLLLAGYDDFNCNVWDAL<br>KADRAGVLAGHDNRVSCLGVTDDGMAVATGSWDSFLKIWN | Expression in Hi5 |
| Bovine Gγ1 | MPVINIEDLTEKDKLKMEVDQLKKEVTLERMLVSKCCEEFRDYVEE<br>RSGEDPLVKGIPEDKNPFKELKGGCVIS | Extraction from Bovine retina |
| Human Gγ2 | MASNNTASIAQARKLVEQLKMEANIDRIKVSKAAADLMAYCEAHAK<br>EDPLLTPVPASENPFREKKFFCAIL | Expression in Hi5 |

**Supplementary Table 3:** Sequences and production sources of JSR1 and G protein subunits.
